# Immunogenicity assessment of PRRS polylactic acid glycolic acid DNA vaccine

**DOI:** 10.1101/650739

**Authors:** Sean Kowalski, John Smith

**Affiliations:** Department of Biochemistry, University of South Carolina, Columbia SC, 29208

## Abstract

In order to enhance the immune effect of DNA vaccine, poly(lactide-co-glycolide (PLGA)] microparticles were prepared by solvent evaporation method, and the porcine reproductive and respiratory syndrome (PRRS) DNA vaccine pCI-ORF5 was adsorbed to the porcine reproductive and respiratory syndrome (PRRS) DNA vaccine. The surface of the microparticles was used to detect the amount of DNA adsorbed by PLGA microparticles, in vitro release, and immunogenicity in mice. The results showed that the DNA adsorption capacity of PLGA particles could reach 0.9% within 6h, and the release in vitro was affected by many factors such as CTAB content, PLGA molecular weight, PLGA concentration and internal water phase volume. After immunizing mice with the naked DNA vaccine, PLGA microparticles were found to significantly enhance the humoral and cellular immunity induced by the adsorbed DNA vaccine, indicating that it has a good application prospect as a vector for delivering DNA vaccines.

## Introduction

DNA vaccines have been used for many years in animal disease prevention. They have the advantages of short preparation period, high specificity, high safety, and can induce cellular immunity and humoral immunity at the same time. However, compared with other vaccines, its immune effect is relatively poor, and multiple, large doses of injection are required, which greatly limits the wide application of DNA vaccines [1-2]. Porcine Reproductive and Respiratory Syndrome (PRRS) is one of the most important infectious diseases that seriously endangers the pig industry and is extremely difficult to effectively control. At present, ORF5 is the major protective antigen gene of Porcine Reproductive and Respiratory Syndrome Virus (PRRSV). The GP5 envelope glycoprotein encoded by ORF5 is an important target protein for inducing neutralizing antibodies and cellular immunity, but constructed based on ORF5. PRRS DNA vaccines often have problems such as low immune titers and low protection. In order to improve the immune effect of the PRRS DNA vaccine, the expression of the target antigen can be enhanced by modifying the vector; secondly, the DNA vaccine can be used in combination with an adjuvant; and third, the delivery mode of the DNA vaccine can be changed. The use of biodegradable high molecular polymers as carriers to deliver DNA vaccines not only helps to increase the immunogenicity of DNA vaccines, but also greatly reduces the amount of DNA vaccine inoculation and the number of inoculations, providing new uses for DNA vaccines. The idea [3]. Polylactic acid-glycolic acid (PLGA) is a biodegradable polymer with good biocompatibility and is widely used in pharmaceuticals, especially as a macromolecular drug delivery carrier [4-5]. The positively charged PLGA/CTAB microparticles prepared by PLGA can effectively adsorb and protect DNA, and the slow release can improve the half-life and immune effect of the adsorbed DNA vaccine in vivo [6]. It has been confirmed in studies of human immunodeficiency virus (HIV) and foot-and-mouth disease virus (FMDV) that PLGA/CTAB microparticles can significantly enhance the immune response induced by DNA vaccines [7-8]. In this study, the positively charged PL-GA/CTAB microparticles were prepared by solvent evaporation method, and then the DNA vaccine was adsorbed onto the surface. Then, the amount of DNA adsorbed by PLGA/CTAB-DNA microparticles and the release in vitro were detected. The immunoenhancing effect of PLGA/CTAB microparticles on the PRRS DNA vaccine after detection in mice will provide an important basis for the development of novel DNA vaccine delivery systems.

## Materials and method

Recombinant PRRSV YA1 strain and PRRSVORF5 gene The eukaryotic expression plasmid pCI-ORF5 was provided by the Animal Virus Laboratory of Huazhong Agricultural University. E. coli strains DH5α, BL21 (DE3), and green monkey kidney cells (Marc-145) were preserved in our laboratory. Prokaryotic expression of PRRSV GP5 protein The recombinant plasmid pKG-GP5 was constructed from this laboratory.

Three kinds of polylactic acid-glycolic acid (PLGA, lactic acid and glycolic acid polymerization ratio of 50:50) with molecular weights of 20, 60 and 100 ku were purchased from Jinan Biotech Co., Ltd.; dichloromethane, cetyltrimethyl Ammonium bromide (CTAB) was purchased from Shanghai Sinopharm Chemical Reagent Co., Ltd.; DMEM, RPMI1640, and calf serum were all products of In-vitrogen. The HRP-labeled goat anti-mouse IgG secondary antibody is a product of Wuhan Aibotek Biotechnology Co., Ltd.

Preparation of PLGA/CTAB microparticles Weigh 0.5 g of PLGA with a molecular weight of 60 ku in a 50 mL centrifuge tube, gently shake it with 10 mL of dichloromethane, completely dissolve the PLGA, and then add 1 mL of PBS to the centrifuge tube (pH 7. 4) Mix the ultra-speed homogenizer for 3 min to form oil/water microspheres. 50 mL of 0.5% CTAB aqueous solution was added dropwise, and mixed again with an ultra-speed homogenizer for 5 min to form water/oil/water microspheres. Finally, the mixture was placed in an open 100 mL three-necked flask and stirred overnight at room temperature on a magnetic stirrer to completely remove methylene chloride. Transfer the mixed solution in the flask to a 50mL centrifuge tube, centrifuge at 10 °C 10000r·min-1 for 10 min, discard the supernatant, wash the pellet twice with sterile ddH2O, freeze-dry in a freeze dryer for 48 h to powder, ie A PLAB/CTAB particle that is positively charged on the surface.

Weigh 10 mg of PLGA/CTAB microparticles into a 1.5 mL centrifuge tube, add 100 μL of PRRSV DNA vaccine pCI-CRF5 (concentration: 1 μg·μL-1), and shake gently for 6 h at 4 °C on a horizontal shaker to allow DNA to be adsorbed to PLGA/ CTAB particle surface. Centrifuge at 10000r·min-1 for 4min at 4 °C, aspirate the supernatant, wash once with 500μL TE, centrifuge at 10000r·min-1 for 10min at 4 °C, absorb the supernatant, and freeze-dry for 36h in a freeze dryer to powder. PLGA/CTAB-DNA microparticles.

Place PLGA/CTAB-DNA microparticles in a 1.5mL centrifuge tube, add 1mL 0.2mol·L-1 NaOH solution, mix well, and let stand at 4 °C for 10h to degrade PLGA and release the adsorbed DNA. 4 °C 10000r • After centrifugation for 10 min at min-1, the supernatant was collected, and the DNA content in the supernatant was measured by a spectrophotometer to calculate the amount of DNA adsorbed by the PLGA/CTAB microparticles (adsorption amount = mass of adsorbed DNA / mass of PLGA microparticles × 100%).

Place the PLGA/CTAB-DNA microparticles in a 1.5 mL centrifuge tube, add 1 mL of PBS buffer to the centrifuge tube, seal with a parafilm, place in a 37 °C incubator, centrifuge the supernatant after 1 d, and pass the split. The photometer was used to detect the amount of DNA released, and a sustained release curve of PL-GA/CTAB-DNA microparticles was prepared. The pellet was resuspended in 1 mL of PBS buffer and placed in a 37 ° C incubator for 30 days.

## Results and discussion

Sixteen copies of 110 μg of plasmid DNA pCI-ORF5 were mixed with 10 mg of PLGA/CTAB microparticles, respectively, and the amount of DNA adsorbed by PLGA/CTAB microparticles was measured at 0, 1, 2, 4, 6, and 8 h after adsorption. The results showed that the adsorption of DNA by PLGA/CTAB microparticles was fast and effective, and the adsorption amount reached 0.5% within 1 h, and then the adsorption rate gradually became slower, reaching a maximum of about 0.9% at 6 h after adsorption (Fig. 1).

**Figure 1.**
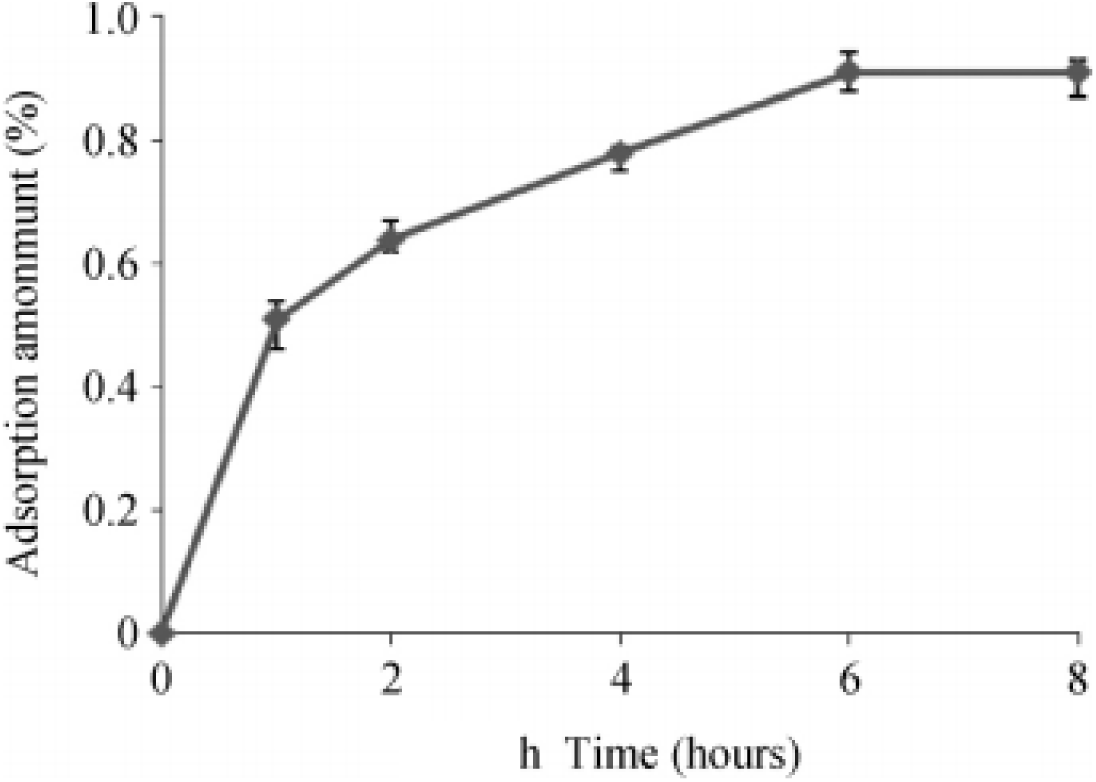
Releasing kinetic of PLGA microparticles.

In order to obtain PLGA/CTAB microparticles with different CTAB contents, PLGA/CTAB microparticles were prepared by washing the precipitates with sterile ddH2O for 1, 2, 3, and 4 times after stirring overnight with magnetic stirrer [9], respectively, after adsorption with plasmid DNApCI-ORF5. The amount of DNA adsorbed and the sustained release were examined. The results showed that with the increase of the number of washings, the content of CTAB decreased gradually, and the amount of DNA adsorbed by PL-GA/CTAB particles washed once or twice was similar to 0.9%, while washing 3 times, 4 The amount of DNA adsorbed by the PLGA/CTAB particles was significantly reduced (Table 1). The PLGA/CTAB microparticles prepared by washing different times were detected by high performance liquid chromatography (HPLC). It was confirmed that the content of CTAB in PLGA/CTAB microparticles decreased with the increase of washing times, indicating the CTAB content in PL-GA/CTAB microparticles. Adsorption of DNA has a large effect.

The PLGA/CTAB microparticles with different CTAB contents were adsorbed to the plasmid DNApCI-ORF5, lyophilized, and PBS was added to measure the amount of DNA released at 37 ° C every 1 d. It can be seen from Fig. 2 that although the CTAB contents of the four PLGA/CTAB-DNA microparticles are different, there is a burst of DNA release for about 1 week. As the number of washings increases, the CTAB content is lower, PLGA/CTAB-DNA. The earlier the bursty release of the DNA of the microparticles, the greater the release and the faster the total release rate.

**Figure 2.**
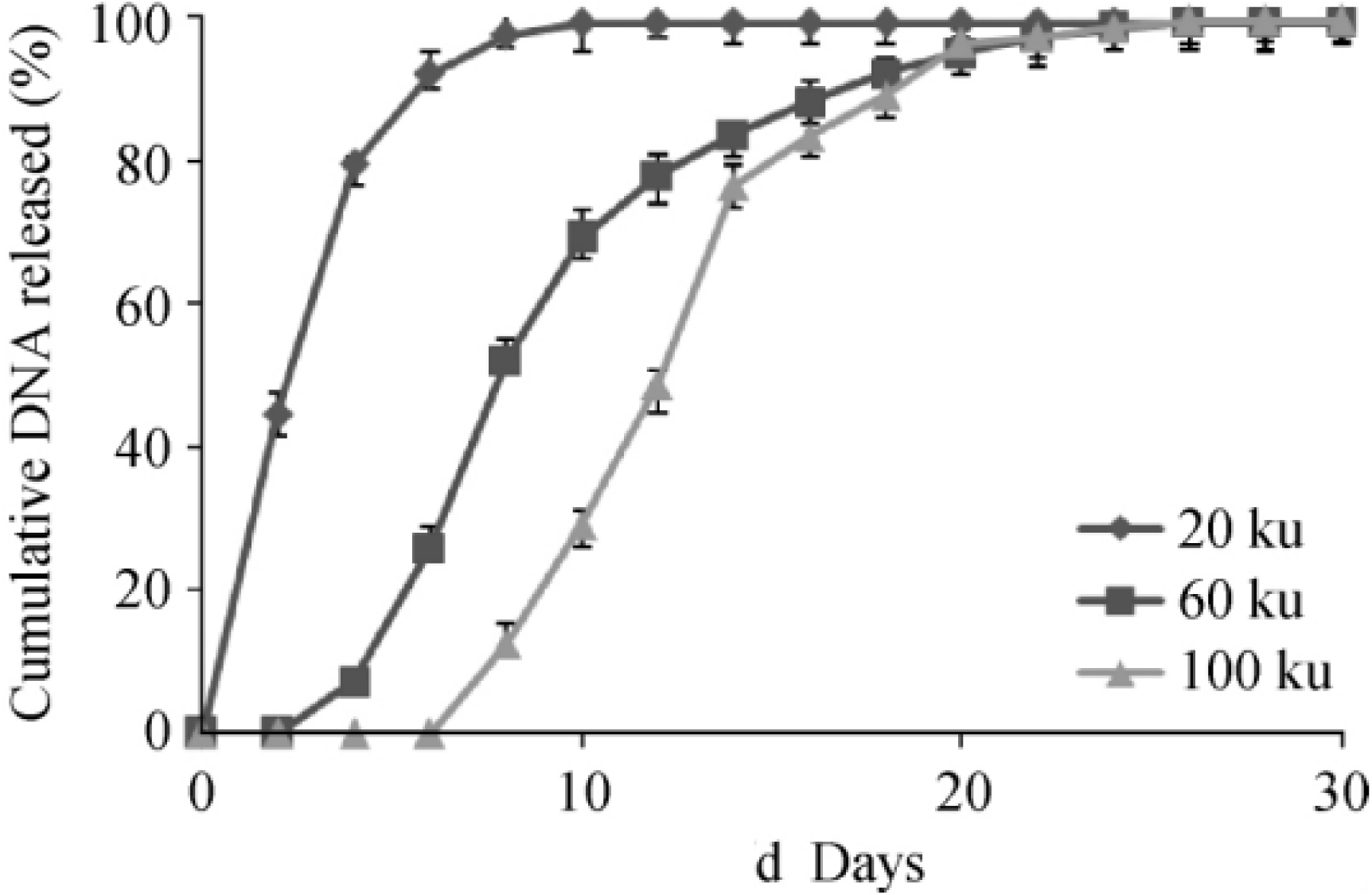
Releasing kinetics of different PLGA microspheres.

PLGA/CTAB microparticles were prepared by using PLGA with molecular weights of 20, 60, and 100 ku, and the DNA was adsorbed, lyophilized, and added to PBS, and the amount of released DNA was measured every 1 day at 37 °C. The results showed that when the molecular weight of PLGA was 20 ku, the DNA was almost completely released in the first week; when the molecular weight was 100 ku, the DNA was released at the second week, and the release was completed in about one week; and the molecular weight was 60 ku on the third day. DNA is released, explosive release for about 2 weeks, and the entire release process is relatively flat (Figure 3).

**Figure 3.**
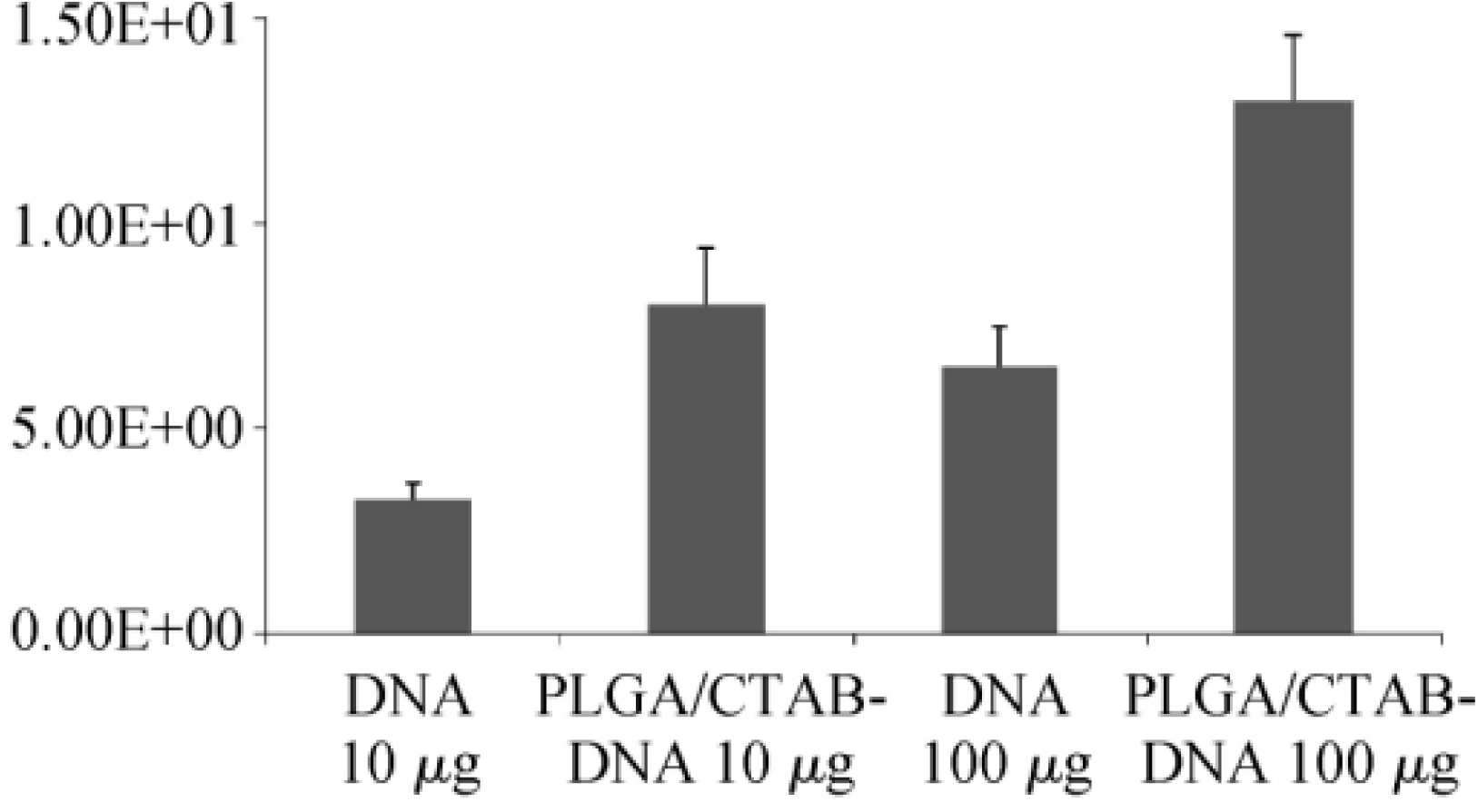
In Vitro luciferase expression.

PLGA/CTAB microparticles were prepared by preparing PLGA into 2%, 5%, and 10% (w/v) dichloromethane solutions. The DNA was adsorbed, lyophilized, and added to PBS. The release was detected at 37 ° C every 1 d. The amount of DNA. The results showed that when the concentration of PLGA was 2% and 10%, the explosive release of DNA by PLGA/CTAB particles was concentrated in the first week and the second week, respectively, and the release amount reached 80% and 70%, respectively. When the concentration was 5%, The explosive release lasted approximately 2 weeks (Figure 4).

**Figure 4.**
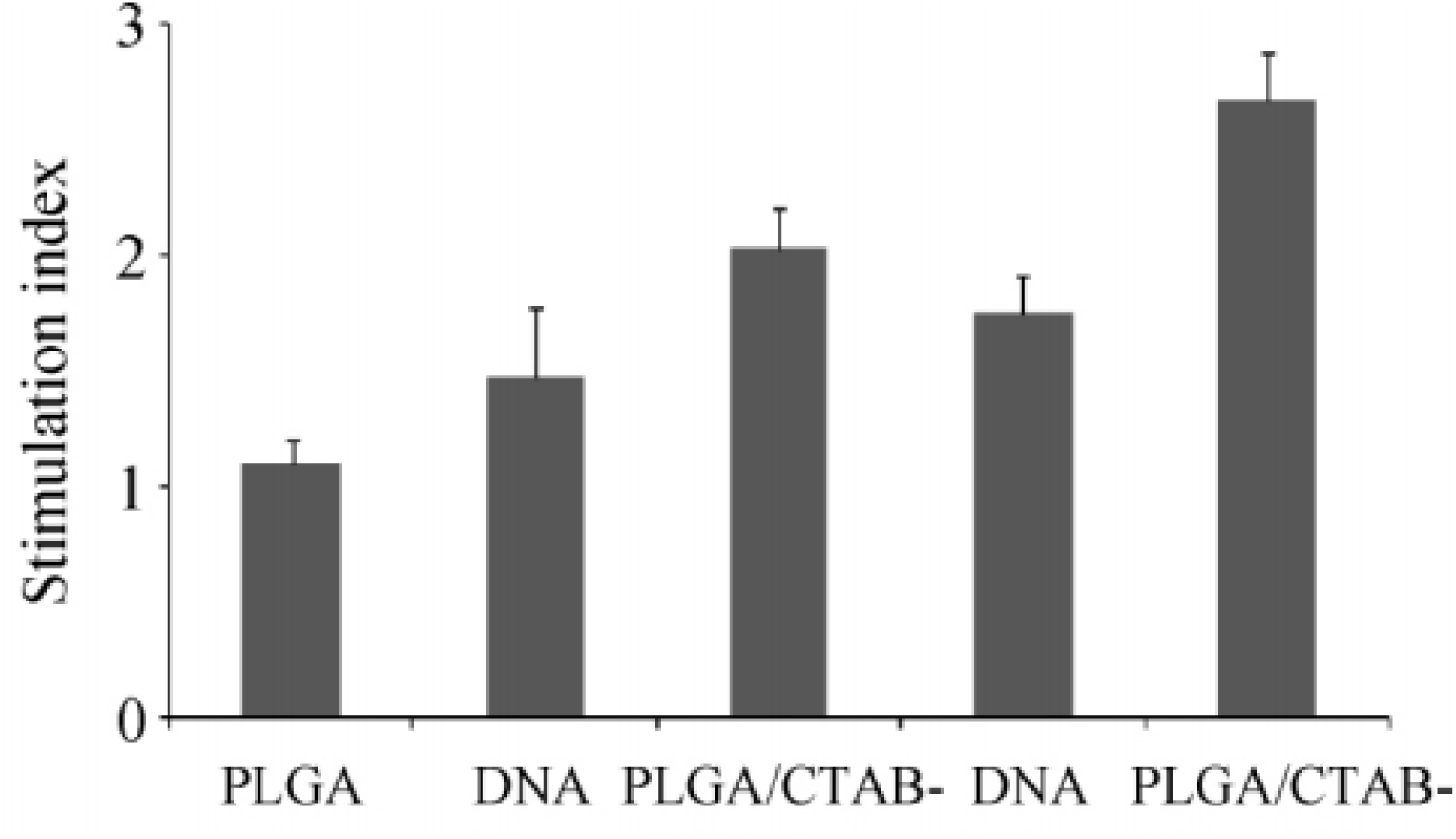
Stimulation index assessment.

PLGA/CTAB microparticles were prepared by ultra-sonic homogenization of PLGA solution with 0.5 mL, 1 mL, and 2 mL PBS, adsorbed with DNA, lyophilized, and added with PBS. The amount of DNA released was measured every 1 d at 37 °C. It was found that when the volume of PBS in the inner aqueous phase was increased to 2 mL, DNA release was detected on the 2nd day, and when it was reduced to 1, 0.5 mL, DNA release was detected on the 4th day, but the release amount was relatively less. As can be seen from Fig. 5, as the volume of the inner aqueous phase is continuously reduced, the release of DNA by PLGA/CTAB-DNA microparticles is gradually slowed down (Fig. 5).

PLGA, naked DNA vaccine pCI-ORF5 and DNA vaccine pCI-ORF5 adsorbed onto PLGA/CTAB microparticles were immunized with mice at different doses, and immunized twice, at intervals of 2 weeks, after the first 2, 4 ELISA antibody levels of anti-GP5 protein in serum were measured at 6, 8 weeks. As can be seen from Figure 6, whether low-dose immunization (10 μg) or high-dose immunization (100 μg), PLGA/CTAB-DNA microparticles induced GP5-specific ELISA antibodies after 4 weeks of first immunization were significantly higher than the same dose of nude DNA vaccines, the difference was extremely significant (P < 0.01), and immunization at low doses (10 μg) induced similar levels of ELISA antibodies to high dose (100 μg) naked DNA vaccines. At the same time, PLGA/CTAB-DNA microparticle-induced GP5-specific ELISA antibody decreased slowly in mice and maintained at a high level for a long time.

The naked DNA vaccine pCI-ORF5 and the DNA vaccine pCI-ORF5 adsorbed on the surface of PLGA/CTAB microparticles were immunized twice at different doses, and the levels of neutralizing antibodies were measured at 8 weeks after the first immunization. As can be seen from Fig. 7, when immunized with the same dose of DNA, the neutralizing antibody of PLGA/CTAB-DNA microparticle immunization group was higher than that of naked DNA vaccine immunization group 8 weeks after the first dose (P<0. 05), and at low dose^1-20^.

The PLGA, the naked DNA vaccine pCI-ORF5 and the DNA vaccine pCI-ORF5 adsorbing the surface of PL-GA/CTAB microparticles were immunized with different doses, and the mice were immunized twice. The spleen lymphocytes of the mice were detected 8 weeks after the first immunization. Specific proliferation. The results showed that when immunized with the same dose of DNA, PLGA/CTAB-DNA microparticle immunization group obtained stronger lymphocyte-specific proliferation 8 weeks after the first immunization, the difference was extremely significant (P<0.01). A DNA vaccine that adsorbs the surface of PL-GA/CTAB particles is able to elicit a stronger cellular immune response (Figure 8).

DNA vaccine is a new type of vaccine developed with the development of modern molecular biology technology. Although it has many advantages, it is difficult to induce a high immune response. In order to achieve the desired immunoprotective effect, multiple, large doses of vaccination are often required. In order to overcome these problems, PLGA/CTAB microparticles were prepared and used as vectors to deliver DNA vaccines. PLGA itself does not adsorb DNA. It must first be combined with CTAB to form a PLGA/CTAB microparticle with a positively charged surface to adsorb DNA. Since DNA is adsorbed on the surface of PLGA microparticles and is not encapsulated inside PLGA microparticles, DNA denaturation and degradation can be avoided to maintain DNA integrity [10]. This study showed that PL-GA/CTAB microparticles can reach 0.5% of DNA adsorption in 1 h in vitro and reach the maximum within 6 h, indicating that this DNA loading method is efficient and rapid. Different PLGA/CTAB microparticles were prepared by changing CTAB content, PLGA molecular weight, PLGA concentration and internal aqueous phase volume, and found that their adsorption and release of DNA in vitro were significantly different. PLGA microparticles prepared with a molecular weight of 60 ku, a PLGA concentration of 5%, an internal aqueous phase PBS volume of 1 mL, and 2 washes of sterile water have a DNA adsorption capacity of 0.9%, and an explosive release amount within 2 weeks in vitro. Nearly 80%, the total release time is more than 3 weeks, the whole release process is longer and more gradual, so it is used to immunize mice to test their immune enhancement effect on PRRSV DNA vaccine. The amount of DNA adsorbed by different PLGA/CTAB microparticles with different CTAB content is mainly due to the difference in the number of charges on the surface, and the diameter has not changed significantly (the diameter is 1∼1.5μm) [11]. A higher amount of DNA adsorption does not produce a higher immune response, and an adsorption amount close to 1% is sufficient to significantly increase the DNA-induced immune response [7]. PLGA/CTAB microparticles can significantly enhance humoral and cellular immunity induced by adsorbed DNA vaccines, on the one hand because they control the slow release of DNA and prevent DNA from being degraded by nucleases in the body; on the other hand, it may be more The DNA vaccine is well delivered to antigen presenting cells (APCs). In vitro transfection experiments with dendritic cells (DC) have shown that PLGA/CTAB microparticles enhance DC capture, gene expression and antigen presentation of DNA vaccines [12]. While DC is the most powerful APC in the immune system, enhancing the antigen presentation of DNA vaccines by DCs will improve the immune response of DNA vaccines. In addition, PLGA/CTAB microparticles are phagocytosed by DC and enter the endosomes. The positive charge on the surface can promote the rupture of endosomes, further enhancing the expression of the adsorbed DNA vaccine in DC, indicating that PLGA/CTAB microparticles have both carrier functions. It may also have the effect of a partial adjuvant, although the mechanism of action of its adjuvant is not fully understood.

## Conclusion

Although PLGA/CTAB microparticles in this experiment can significantly enhance the immune response induced by the adsorbed PRRS DNA vaccine, it still does not reach the level of inactivated or attenuated PRRSV vaccine. It may be because this experiment only uses the single gene ORF5 to immunize, but the ORF5 gene alone. The constructed DNA vaccine is often difficult to stimulate the body to produce a higher immune response [13]. In the subsequent trials, DNA vaccines expressing the ORF5 gene and the ORF6 gene will be constructed separately, and then PLGA/CTAB microparticles will be loaded simultaneously to further enhance the body’s immunity to PRRSV. It has been confirmed that PLGA/CTAB microparticles that adsorb one or more plasmid DNAs can induce a stronger immune response than the naked DNA vaccine, providing a more comprehensive protection for the organism [12], which is undoubtedly PLGA/CTAB. The use of microparticles as a DNA vaccine vector provides a broader perspective.

